# Phosphorylation at serine 214 correlates with tau seeding activity in an age-dependent manner in two mouse models for tauopathies and is required for tau transsynaptic propagation

**DOI:** 10.1101/2024.07.22.604618

**Authors:** Pablo Martinez, Nur Jury-Garfe, Henika Patel, Yanwen You, Abigail Perkins, Yingjan You, Audrey Lee-Gosselin, Ruben Vidal, Cristian A. Lasagna-Reeves

## Abstract

Pathological aggregation and propagation of hyperphosphorylated and aberrant forms of tau are critical features of the clinical progression of Alzheimer’s disease and other tauopathies. To better understand the correlation between these pathological tau species and disease progression, we profiled the temporal progression of tau seeding activity and the levels of various phospho- and conformational tau species in the brains of two mouse models of human tauopathies. Our findings indicate that tau seeding is an early event that occurs well before the appearance of AT8-positive NFT. Specifically, we observed that tau phosphorylation in serine 214 (pTau-Ser214) positively correlates to tau seeding activity during disease progression in both mouse models. Furthermore, we found that the histopathology of pTau-Ser214 appears much earlier and has a distinct pattern and compartmentalization compared to the pathology of AT8, demonstrating the diversity of tau species within the same region of the brain. Importantly, we also observed that preventing the phosphorylation of tau at Ser214 significantly decreases tau propagation in mouse primary neurons, and seeding activity in a *Drosophila* model of tauopathy, suggesting a role for this tau phosphorylation in spreading pathological forms of tau. Together, these results suggest that the diverse spectrum of soluble pathological tau species could be responsible for the distinct pathological properties of tau and that it is critical to dissect the nature of the tau seed in the context of disease progression.

## INTRODUCTION

The microtubule-associated protein tau is required for microtubule assembly, axonal transport, and neurite outgrowth. Most of the biological functions of tau are modulated by site-specific phosphorylations^1^. Tau is encoded by a single gene, with 6 splice isoforms, ranging in size from 352 to 441 amino acids, which are expressed in the human CNS^2,3^. These isoforms differ from each other by the presence of 0, 1, or 2 N-terminal inserts and 3 or 4 tandemly arranged microtubule-binding repeats. Tau undergoes many post-translational modifications including glycosylation, ubiquitination, glycation, polyamination, nitrosylation, and truncation. Nevertheless, the most important and disease-relevant tau post-translational modification is hyperphosphorylations^4^, which alters tau’s biological functions and causes tau self-assembly and accumulation as neurofibrillary tangles (NFT), a hallmark of Alzheimer’s Disease (AD) and other neurodegenerative diseases^5,6^. All tau isoforms contain at least 30 phosphorylation sites^4^, most of which are believed to be unphosphorylated under physiological conditions. However, under pathological conditions, hyperphosphorylation of various sites occurs resulting in tau accumulation^7,8^. Tau phosphorylation, especially at specific sites, reduces its affinity for microtubules^9,10^. Therefore, it is not surprising that considerable attention has been paid to determining which protein kinases and phosphatases control tau phosphorylation^11^.

In AD, several studies have described a spatial and temporal pattern in the appearance of tau tangles in patient brains that follow neuronal networks and correlate with cognitive decline^12^. In the case of AD, tau tangles first appear in the transentorhinal region and progress along anatomical pathways to the hippocampus and eventually the neocortex^12^. Similar temporal progression of tau pathology is observed in argyrophilic grain disease (AGD), though the brain regions involved differ^13,14^. Likewise, the spatial distribution of tangles is distinct in other tauopathies^15^. While still a contentious topic, strong evidence supports the idea that propagation of pathological tau species occurs between cells^16–18^, which suggests that this could contribute to the spatial and temporal pattern of the tangles observed in different tauopathies. Cell culture studies have demonstrated that misfolded tau aggregates can mediate a template misfolding or “seeding” of normal, monomeric tau to induce intracellular tau aggregation^19–21^. Furthermore, in vivo tau spreading models support the mechanism of tau propagation. Injection of recombinant tau aggregates or brain lysate containing tau aggregates into the brains of WT or young transgenic mice can induce robust pathology at the site of injection and in anatomically connected regions^20,22,23^. The induction of tau pathology in mice that do not otherwise develop tau inclusions supports the concept of seeding and the propagation of tau aggregates to anatomically connected neurons.

Blocking the spread of the tau seed with antibodies has been proposed as a viable therapeutic approach. However, the facts that the tau seed structure remains unknown, and recent studies indicating a distinct molecular diversity of the seeding-competent tau in in vitro and in vivo models^23–27^, make it extremely challenging to determine the exact epitope or post-translational modification to target. Recently, studies suggested that tau antibodies directed to the middle region neutralized tau propagation more efficiently than antibodies targeting the N-terminal region^28^. The ability of many of these antibodies to block tau seeding has been tested in vitro using brain tissue extract from a low number of donors at a single point of the pathology^29,30^. Considering the heterogeneity of the tau seed among individuals with tauopathies and even in disease progression^31^, the efficiency of each of these antibodies to inhibit tau seeding could be diminished if tested in a different experimental model or even at a different time point in humans. Therefore, it is necessary to understand the nature of the tau species involved in spreading and the precise seeding/template.

In the present study, we utilized a well-characterized FRET-based biosensor cell line^32^ to profile the temporal progression of tau seeding activity in the brains of two mouse models of human tauopathies^33,34^. First, we show that tau seeding activity is a pathological event that happens several weeks earlier than the appearance of the first symptoms in these mouse models. We then profiled the seeding activity at different ages from these mouse models to the levels of a diverse set of phospho- and conformational tau species analyzed from the same brain samples. We found that the level of phosphorylated tau at serine 214 (pTau-Ser214) correlated with tau seeding activity within the individual disease progression in both mouse models for tauopathies. We also observed that tau seeding occurred before the appearance of conventional AT8-positive NFTs and that pTau-Ser214 histopathology manifested earlier and displayed a distinct pattern and compartmentalization compared to AT8 pathology, demonstrating the diversity of tau species within the same brain region. Furthermore, preventing the phosphorylation of tau at Ser214 by introducing the amino acid change from serine to alanine significantly reduced tau propagation in mouse primary neurons, and interestingly reduced seeding activity in a *Drosophila* model of tauopathy, suggesting a role for this tau phosphorylation in the spreading and formation of toxic aberrant forms of tau. Our study highlights the importance of dissecting the nature of the tau seed both in the context of individual disease progression and in different tau models that will allow us to better dissect the mechanism behind the formation of distinct tau species at different stages of disease progression.

## MATERIALS AND METHODS

### Transgenic mouse model

Mice were housed at the Indiana University School of Medicine (IUSM) animal care facility and were maintained according to USDA standards (12-hr light/dark cycle, food, and water *ad libitum*), following the Guide for the Care and Use of Laboratory Animals (National Institutes of Health, Bethesda, MD). The PS19 mouse model^33^, which overexpresses human 1N4R tau with the P301S mutation, was directly purchased from Jackson Laboratories (stock number 008169). A second mouse model that overexpresses human 0N4R tau with the P301S mutation was developed and provided by Dr. Ruben Vidal^34^. Animals were anesthetized and euthanized according to IUSM Institutional Animal Care and Use Committee-approved procedures. For described experiments 2, 3, 4, 5, 6, and 9-month-old animals were utilized. Mice were perfused transcardially with PBS before decapitation. Brains were extracted and the right hemisphere was stored at −80 for biochemical analysis and the left hemisphere was formalin fixed for the preparation of paraffin blocks as previously described^35^.

### Mouse brain samples preparation and immunoblot analysis

Both P301S (0N4R) and P301S (1N4R) mouse brains were homogenized at a 1:10 (w/vol) ratio of the brain in 1X TBS supplemented with protease inhibitor cocktail (Roche), using a Bead Beater (1 min) and then sonicated for 1 min (50% amplitude, pulse on 12 sec, pulse off 3 sec). Samples were then centrifuged at 15,000 rpm for 10 min at 4 °C. TBS soluble supernatant was aliquoted into fresh tubes. For immunoblot analysis, TBS soluble fractions were run on NuPAGE 4-12% Bis-Tris protein gel (Invitrogen) and transferred to a nitrocellulose membrane. The primary antibodies used against tau species were PHF1 (1:1000, from Peter Davies), MC1 (1:1000, from Peter Davies), anti-pS214 tau (1:1000; Abcam ab170892), anti-pT217 tau (1:1000; ThermoFisher, 44-744), total human tau HT7 (1:1000; ThermoFisher, MN1000), anti-tau pS202/T205 (AT8; 1:1000; Invitrogen, MN 1020), anti-tau pT212 (1:1000; Invitrogen, 44-740G), anti-tau pT212/S214 (AT100; 1:1000; Invitrogen, MN1060), anti-tau pT231 (PHF6; 1:1000; EMD Millipore, MAB5450), anti-tau pS262 (1:1000; Abcam, ab131354) and anti-Vinculin (1:1000; Sigma, V9131). The secondary antibodies used were goat anti-mouse HRP IgG (1:5000; Invitrogen, A16066) and goat anti-rabbit HRP IgG (1:5000; Invitrogen A16066). For each analysis, the relative level of each tau antibody and Vinculin was quantified from western blot images using ImageJ software (NIH, v.1.53t). The level of each tau species was normalized to HT7 to determine the relative abundance of each tau species with respect to total tau levels.

### Flow cytometry and analysis of seeding activity

The seeding assay was performed as previously described^35,36^. Tau RD P301S FRET Biosensor cells were plated at 35,000 cells in 130 µl media per well in a 96-well plate, then incubated at 37 overnight. The next day, cells were transfected with brain lysate at a concentration of 20 µg total protein per well using Lipofectamine 2000 and incubated at 37 for 2 days. Cells were harvested by trypsinization and fixed with 2% PFA for flow cytometry. Flow Cytometry was conducted on a BD LSRFortessa™ X-20 with a High Throughput Sampler. BV421 channel was used to detect CFP (405nm Ex, 450/50 Em filter), and BV510 channel to detect FRET signal (405nm Ex, 525/50 + 505LP Em filter), with compensation used to remove the CFP emission from the FRET signal. Gating cells with a positive FRET signal, seeding was quantified by measuring % positive cells and median fluorescent intensity (MFI) of the compensated BV510 channel. Integrated FRET density was calculated by multiplying % positive cells and MFI.

### Correlation Analysis

For each case, the relative level of each tau antibody and Vinculin was quantified from western blot images as described ^35^. The level of each tau species was normalized by Vinculin, then normalized again to total human tau (HT7 normalized by Vinculin) to give a relative abundance for each tau species. Spearman’s rank correlation between % positive FRET signal and tau species abundance was used to identify tau species that correlate with seeding. For graphing of the results, each species’ abundance was normalized to the average abundance over all cases. This ensures that the average of normalized abundance is equal to 1 for each tau species, to enable comparison of each species on the same axis range. Since all abundances are relative, this final normalization step does not affect the correlation analysis.

### Brain Section Immunofluorescence

Paraffin sections from 9-month-old mice were deparaffinized, rehydrated, and washed in 0.01 M PBS 1X plus Triton X-100 0.01% three times for 5 min each. After blocking in normal goat serum for 1 hr, sections were incubated overnight with AT8 antibody (1:200) and anti-pTau-Ser214 antibody (1:200). The next day, the sections were washed in PBS 1X 3 times for 10 min each and then incubated with goat anti-mouse Alexa Fluor 488 (1:700, Invitrogen,) and goat anti-rabbit Alexa Fluor 568 (1:700, Invitrogen,) for 1 hr. Sections were washed and mounted in Fluoromount (Sigma, F4680) mounting medium with DAPI (Vector Laboratories). The sections were examined using a Nikon A1-R laser scanning confocal microscope coupled with Nikon AR software (v.5.21.03).

### Brain sections Immunohistochemistry (IHC)

Immunohistochemistry was performed on paraffin-embedded sections. In brief, sections (5 μm) were deparaffinized and rehydrated. After blocking in normal goat serum for 1 hr, sections were incubated overnight with AT8 antibody (1:100) or anti-pTau-Ser214 antibody (1:100). The next day, the sections were washed in PBS 1X three times for 10 min each and then incubated with biotinylated goat anti-mouse IgG or goat anti-rabbit IgG and visualized using an ABC reagent kit (Vector Laboratories) according to the manufacturer’s recommendations. Bright-field images were acquired using a Leica Application Suite X 3.6.0.20104 on a Leica DMi8 microscope. Sections were counterstained with hematoxylin (Vector Laboratories) for nuclear staining.

### Synaptosomes isolation

Synaptic and cytosolic fractions were isolated using the Syn-PERTM synaptic protein extraction reagent (Thermo Scientific, 87793) using the manufacturer’s instructions. Briefly, hippocampal tissue was homogenized with Syn-PER reagent and the PierceTM protease/phosphatase inhibitor cocktail (Thermo Scientific, A32959,). Then, samples were centrifugated at 1200 G for 15 min, and the remaining supernatant was centrifuged at 15 000 G for 20 min to obtain synaptosome pellet and cytosolic supernatant. Ten micrograms of each sample were heated at 95°C for10 min in Laemmli sample buffer and subjected to SDS-PAGE for 1 h min at 100 V. Proteins were wet-transferred to a nitrocellulose membrane for 2 h at 400 mA. Membranes were blocked with EveryBlot blocking buffer (Bio-Rad, 12010020), followed by blotting protocol, as previously mentioned.

### Mouse primary cortical neurons

Primary cortical neurons were obtained from mouse postnatal animals (P0). Briefly, P0 mouse pups were obtained following ethical approval procedures. Pups were rapidly decapitated, and the brains were carefully removed and placed in cold PBS 1X. Under a dissecting microscope, the cerebral cortices were isolated from the meninges and hippocampi. Isolated cortices were mechanically dissociated into small pieces using sterile scalpels. The tissue fragments were then incubated for 15 minutes at 37°C in an enzyme solution containing trypsin (Thermo Fisher, 15090046) 2.5% (wt/vol) in PBS 1X supplemented with 20 U/ml DNAse I (Sigma-Aldrich, 11284932001). Following incubation, the tissue was gently triturated using fire-polished Pasteur pipettes of varying diameters to achieve a homogenous cell suspension. The cell suspension was centrifuged at 200 x g for 5 minutes at room temperature. The supernatant was discarded, and the cell pellet was resuspended in a complete neural culture medium containing Neurobasal medium (Invitrogen, 21103049) supplemented with 2% B27 supplement (Invitrogen, 17504044), 1% Penicillin/Streptomycin (Invitrogen, 15140148), and 1X of 200 mM of Glutamax (Invitrogen, 35050061). Cell viability was assessed using trypan blue, and the cells were plated at a density of 50.000 on poly-D-lysine-coated (Invitrogen, A3890401) microfluidic chambers (Xona Microfluidics, SND450). Cultures were maintained in a humidified incubator at 37°C with 5% CO2. After 2 hours, half of the medium was replaced with fresh neuronal media. Thereafter, half of the medium was exchanged with fresh neuronal media every other day.

### Microfluidic system

Microfluidic chambers were used from Xona, No. SND450. The sterile chambers were mounted on glass coverslips previously treated with poly-L-lysine for 30 min at 37. Once adhered, chambers were incubated at 80 for 1 hr to allow the adhesion between the chamber and the glass. Chambers were exposed to UV light for 5 min for sterilization followed by the application of 100 μl of medium to each chamber, and then 50,000 primary WT mouse neurons were plated (Adapted from previous work^37^).

### Adeno Associated Virus (AAV) production and transduction

P301S (0N4R) was cloned and packaged in an AAV9 vector as described before^38^, with modifications. Briefly, V5-tagged Tau (1N4R^P301L^) and Tau (1N4R^P301L/S214A^), and GFP expression plasmids were cloned into an AAV vector and then the different AAV were packaged by VectorBuilder.

For transduction in microfluidics, 1µL with 10^10^ viral particles was applied directly to the upper-left compartment and allowed to diffuse through the left chamber by volume difference. Maintenance media were changed every 48 hours.

### Drosophila stocks and genetics

Drosophila melanogaster stocks and crosses were maintained on Nutri-fly Bloomington formulation (Genesee Scientific) at 25°C in a 12-hour light/dark cycle. All the fly experiments were performed on day 10 post-eclosion. Transgene overexpression was achieved with the Gal4/UAS system, using the GMR-Gal4 line (BDSC, #9146). For human tau overexpression, tau 2N4R harboring the P301L and S214A mutation sequences were cloned downstream of the Gal4-responsive upstream activating sequences into the pUAST plasmid (Vectorbuilder) and then microinjected in fly embryos (Bestgene). Soluble fly head homogenates were obtained by manual homogenization of at least 10 fly heads in TBS 1X and proteinase/phosphatase cocktail inhibitors (Thermo) followed by centrifugation at 15,000 rpm for 10 min. Supernatants were stored at −80 °C.

### Statistical Analyses

All experimental analyses and data collection was performed in a blinded fashion. All p-values were determined using the appropriate statistical method using GraphPad Prism, as described throughout the manuscript. For simple comparisons, a Student’s t-test was used. For multiple comparisons, ANOVA followed by the appropriate post hoc analysis was utilized. Data are presented as the mean ± SEM (*p < 0.05, **p < 0.01, and ***p < 0.001). To identify correlations between phospho-tau epitopes and seeding activity Spearman’s correlation was used.

## RESULTS

### Tau seeding activity is an early pathological manifestation and increases with age in two mouse models of tauopathy

Considering the significant heterogeneity of the tau seed observed among individuals with tauopathies^31^ as well as in disease progression, we aimed to investigate and characterize the tau seeding activity in relation to disease progression. To achieve this, we used two different mouse models of human tauopathies, for a comprehensive understanding of the underlying mechanisms of tau pathology. We chose these models due to their relevance and proven efficacy in studying the pathological changes in tau protein ^33,34^. One of the models expresses the human tau isoform 1N4R with the P301S mutation (P301S (1N4R), also known as the PS19 mouse model)^33^. This model has been extensively characterized and is known to develop NFTs-like pathology at 6 months of age and behavioral impairment at 9 months of age. The second model expresses the smallest 4-repeat human isoform of tau (0N4R) with the P301S mutation (P301S (0N4R))^34^. In this model, it is possible to observe AT8-positive tau deposits as early as 5 to 6 months of age, and, starting at 8 months of age, these mice show motor and behavioral deficits such as the inability to spread hind limbs upon tail elevation, lack of grooming, inability to grip a wire cage, and hind limb dystonia^34^. To measure tau seeding bioactivity in these models, we utilized a widely used cell-based assay that relies on flow cytometry detection of FRET^32^. FRET can be created by the intracellular aggregation of the endogenously expressed repeat domain of tau (TauRD) fused with the cyan fluorescent protein (TauRD-CFP) and the yellow fluorescent protein (TauRD-YFP) in stably expressing HEK-293 cells^32^. We collected mouse brains aged 2-9 months from both tauopathy models. Each brain was homogenized in TBS buffer with no detergent, and lysates were transfected into the tau-biosensor cell line. After forty-eight hours, we detected strong seeding activity from both tauopathy model brain homogenates, whereas WT and tau knock-out mice did not display any seeding activity (Fig. 1). The seeding activity was observed as early as in 2-month-old brain samples and seems to be increased over time in both tau models. This data supports the notion that tau seeding is an early and robust marker of pathology progression and occurs much earlier than the appearance of standard NFT histopathology in two different mouse models of human tauopathies.

**FIGURE 1.**
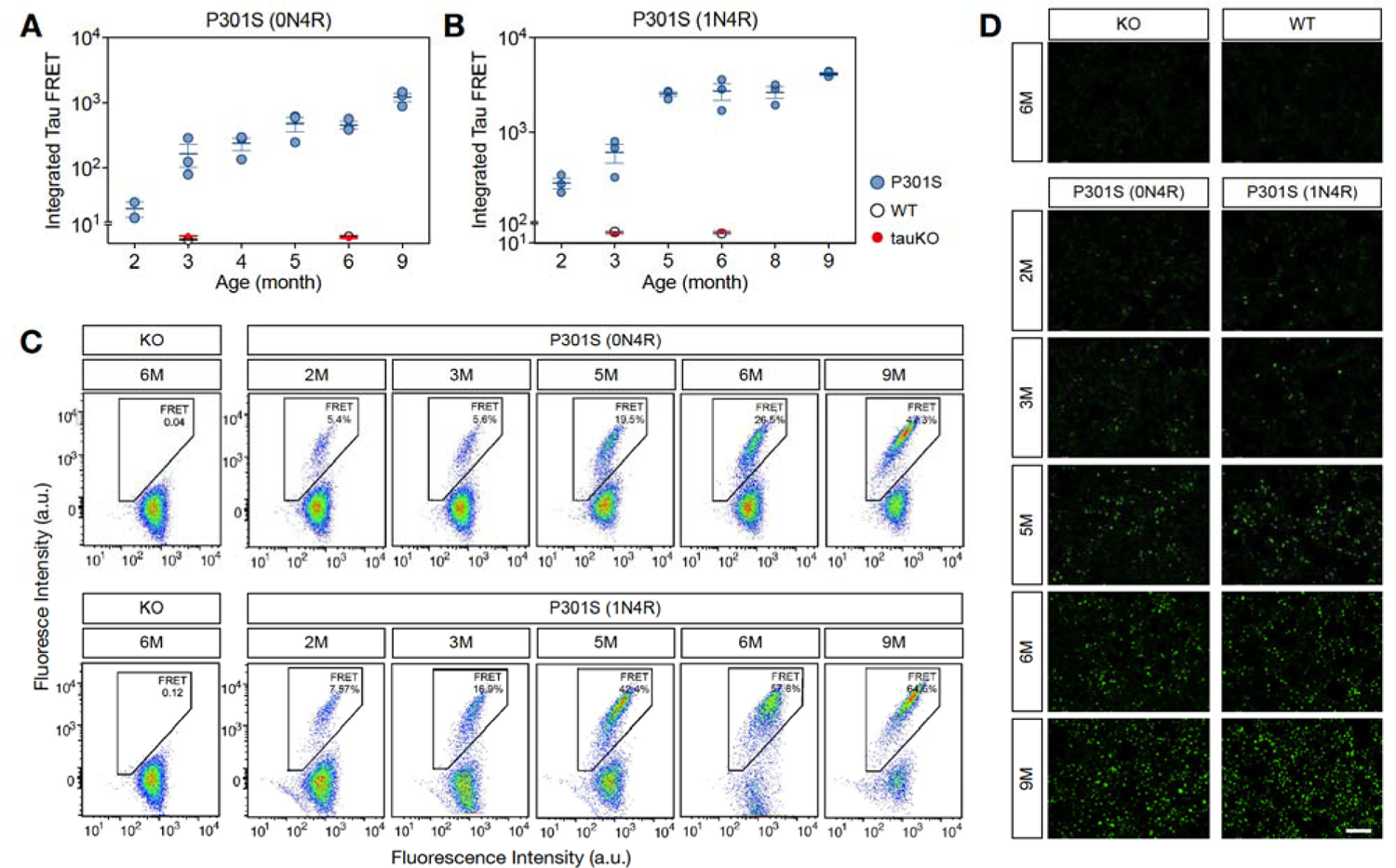
Tau seeding activity is an early pathological manifestation and increases with age in two models of tauopathy. (**A, B**) Integrated FRET quantification from cells transfected with TBS-based brain lysates from 2-9 months old P301S (0N4R) (**A**) and P301S (1N4R) (**B**) mouse models. Each dot represents the average of three technical replicates per lysate. (**C**, **D**) representative integrated FRET signals (**C**) and fluorescent images (**D**) from HEK tau-biosensor cell lines used from tau seeding activity qualifications. Experiments were performed with an n=3 using technical triplicates for each n. Values are represented as the mean ± s.e.m.; n=3. Scale bar: 100 µm.

### Tau phosphorylation at serine 214 correlates with seeding activity in disease progression in two mouse models of tauopathy

After characterizing the temporal profile of tau seeding activity in the brains of both tauopathy mouse models, we aimed to correlate these profiles with a diverse set of phospho-tau and conformational tau species. To do so, we utilized the same mouse brain TBS-lysates used for seeding activity analysis and performed western blot analysis using the following antibodies: HT7 for total human tau, AT8 for pTau-S202/T205, AT100 for pTau-T212/S214, PHF6 for pTau-T231, anti pTau-T212, anti pTau-S214, anti pTau-T217, anti pTau-S262, PHF1 for pTau-S396/S404 and MC1 that recognizes amino acids on both the N-terminal (amino acids 7-9) and within the repeat domain (amino acids 313-322) of tau (Fig. 2A and 2B). Spearman’s rank correlation between the percentage of positive FRET signal and tau species abundance was used to identify tau species that correlate with the temporal evolution of tau seeding. To enable a comparison of each tau species on the same axis (Fig, 2C and 2D), each species’ abundance was normalized to the average abundance over all the cases. This ensures that the average in normalized abundance is equal to 1 for each tau species. In this study, we focused on tau species with a positive correlation with seeding activity, which allowed us to determine the species that followed the disease progression in the context of the amount of seeding activity. In the case of the P301S (0N4R) mouse model, pTau-Ser214 species displayed a significant positive correlation (p=0.007) with tau seeding activity in overall disease progression (Fig. 2C). Furthermore, the tau species with a significant positive correlation with the tau seeding activity in the P301S (1N4R) model were pTau-Thr217 (p=0.0226) and pTau-Ser214 (p=0.0142) (Fig. 2D).

**FIGURE 2.**
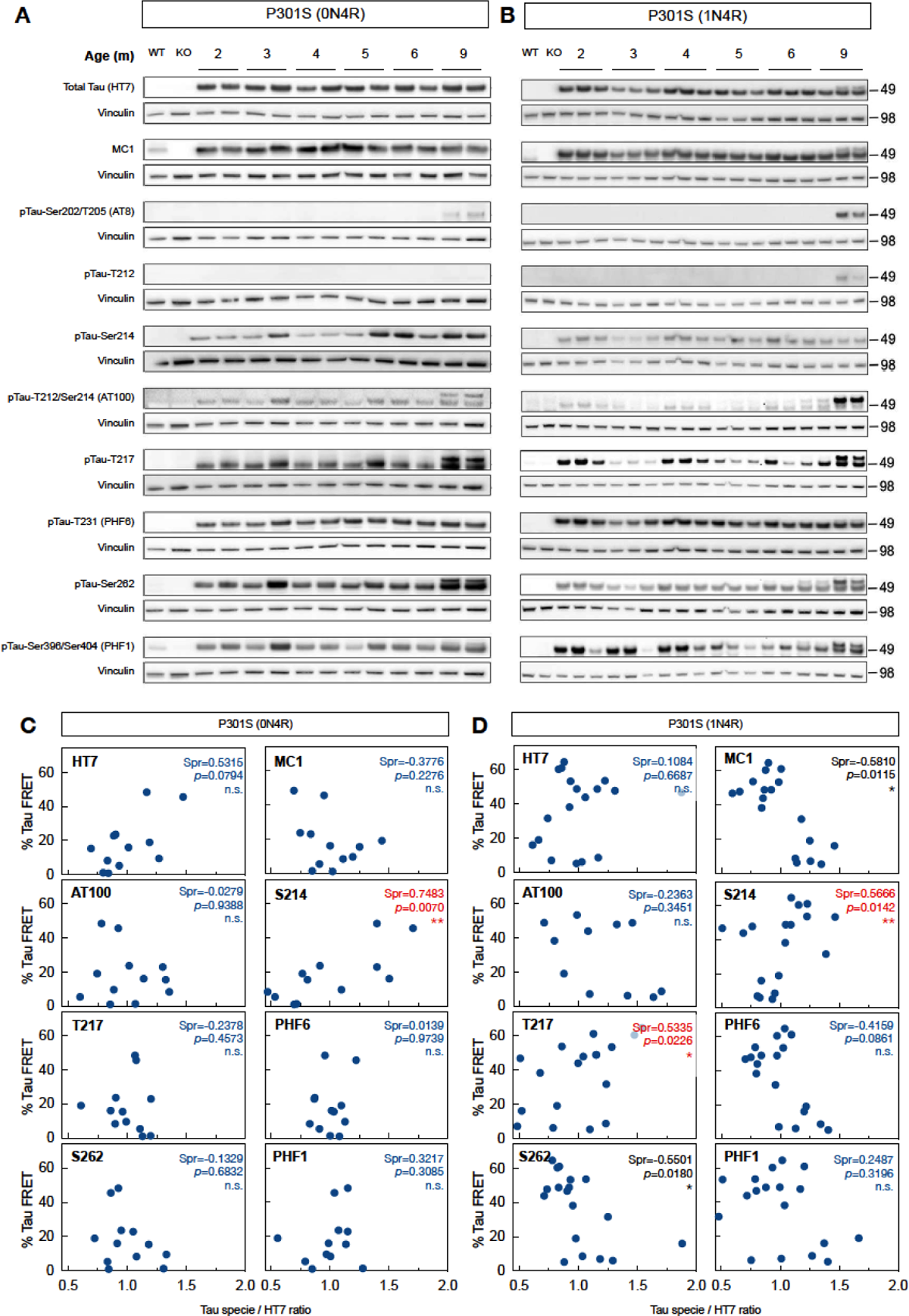
Tau phosphorylation at serine 214 correlates with seeding activity in disease progression in two mouse models of tauopathy. (**A, B**) Characterization of different tau species (HT7 for total human tau, AT8 for pTau-Ser202/Thr205, AT100 for pTau-Thr212/Ser214, PHF6 for pTau-Thr231, anti-pTau-T212, anti-pTau-Sser214, anti-pTau-Thr217, anti-pTau-Ser262, PHF1 for pTau-Ser396/S404, and MC1 that recognizes misfolded tau) in full brain TBS-lysates from 2-9 months P301S (0N4R) (**A**) and P301S (1N4R) (**B**) mouse models. (**C, D**) Correlation between percentage of FRET signal in the P301S (0N4R) (**C**) and P301S (1N4R) (**D**) mouse models with phosphorylated tau species. Dots indicate the average of three animals, each one quantified in technical triplicates. Red values indicate significance for positive correlations. Values are represented as the mean ± s.e.m.; n=3.

Considering that pTau-Ser214 was the only tau species with a significant positive correlation with tau seeding activity during disease progression in both mouse models of tauopathy, ruling out any specific model effect, these results suggested that tau phosphorylation at serine 214 could be a reliable marker to correlate and/or follow the pathogenic evolution of tau species with seeding activity despite the heterogeneity of the tau seed among individuals and animal models of tauopathy^31^.

### Tau phosphorylated at serine 214 is observed at an early-stage disease with distinct histopathological features from AT8-positive tau species

To characterize the histopathological deposition of pTau-Ser214 during disease progression, hippocampal sections from both tauopathy models were stained with anti pTau-Ser214 antibodies at three different ages (3, 6, and 9 months). Interestingly, pTau-Ser214 pathology was observed in the hippocampus of 3-month-old P301S (0N4R) and P301S (1N4R) mice (Fig. 3A and 3B), much earlier than the appearance of NFT-like pathology ^33,34^. In both models, pTau-Ser124 pathology was initially observed in the CA3 region of the hippocampus and mossy fiber projecting from the dentate gyrus. An increase of pTau-Ser214 staining was observed in both models at 6 months of age in the dentate gyrus, CA3, and other areas of the hippocampus such as CA2 and even CA1 in the case of the P301S (0N4R) model (Fig. 3A and 3B, top panels). At 9 months of age, which represents a late stage of the pathology for both models^33,34^, pTau-Ser214 staining was significantly increased in the hippocampus, mossy fibers, and dentate gyrus (Fig. 3A and 3B, top panels). The monoclonal antibody AT8, which recognizes the double phosphorylation at serine 202 and Threonine 205 on aggregated tau protein, has been widely recognized as a gold standard for the detection of tau pathology, particularly in the form of NFTs by IHC^12,39^. Therefore, we stained adjacent sections from both models with the AT8 antibody. As previously reported for both mouse models used in this study, AT8-positive tau was observed at later stages of pathology (Fig. 3A and 3B, bottom panels). In the P301S (0N4R) model, AT8 staining was barely observed at 6 months with a substantial increase in signal at 9 months (Fig. 3A, bottom panel). Similarly, in the P301S (0N4R) model, AT8-positive tau pathology was observed at 9 months (Fig. 3B, bottom panel).

**FIGURE 3.**
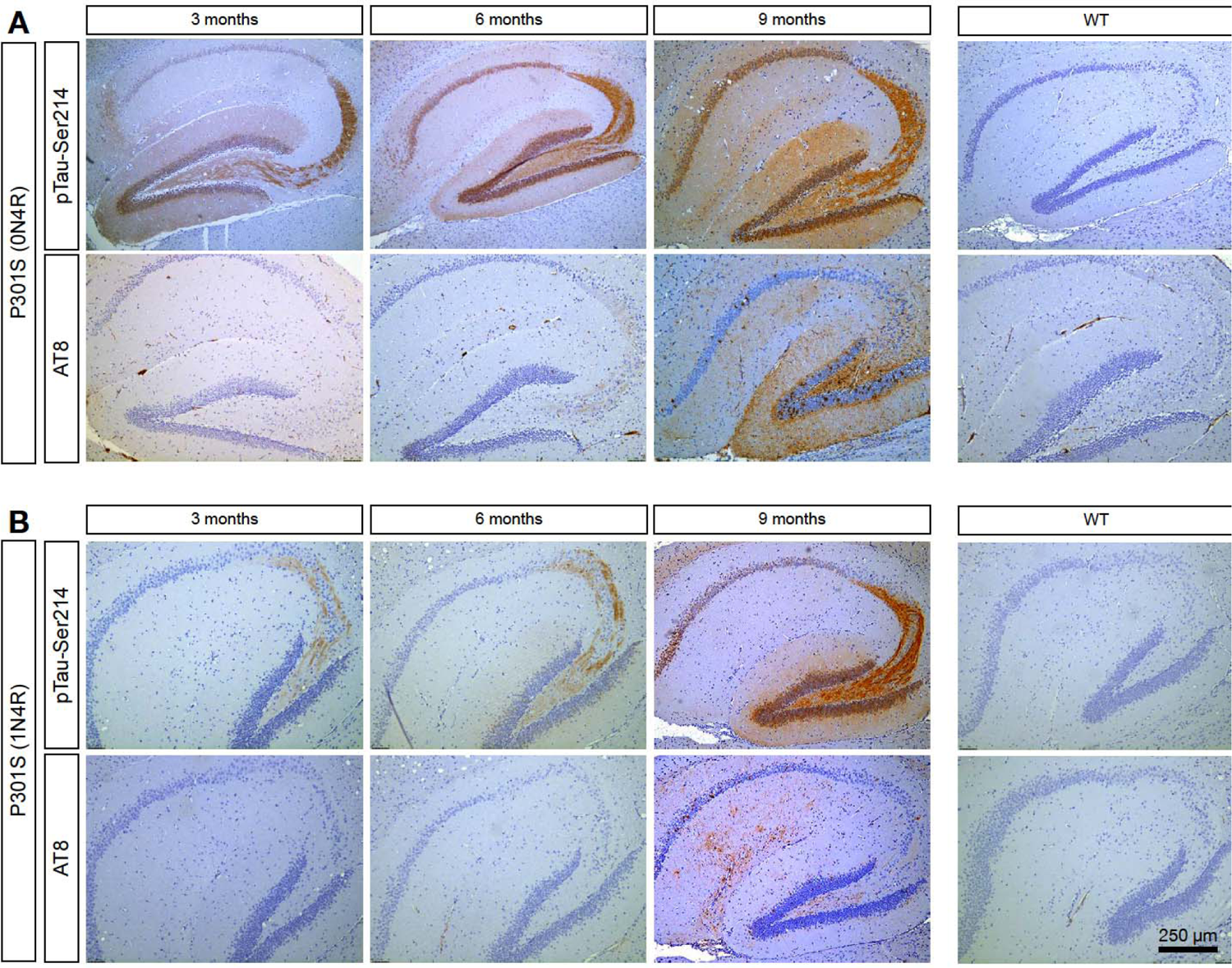
Tau phosphorylated at serine 214 is observed at an early stage of the disease and develops before the formation of AT8-positive tau inclusions. Sagittal hippocampal brain sections from both (**A**) Tau P301S (0N4R) and (**B**) Tau P301S (1N4R) tauopathy models were stained using specific antibodies against pTau-Ser214 and AT8 at 3, 6, and 9-month-old to characterize the histopathological deposition of these phospho-tau species. No positive immunostaining was observed in wild-type mice (right panel). Staining was performed with an n=3.

To better characterize the differences observed at a late stage of pathology between AT8 and pTau-Ser124 pathology, we performed IHC of both markers in 9-month-old mice from both tauopathy models. Upon closer examination of the hippocampus of brain sections using higher magnification, pTau-Ser214 pathology was widely distributed in both tau models. The staining was not limited to the cytoplasm but was predominantly distributed to the axons in the CA3 and DG regions (Fig. 4, top panel). Whereas in adjacent sections AT8 staining revealed the presence of NFTs-like pathology, which is not detected by the pTau-Ser214 antibody, in the hippocampus and DG from both mouse models (Fig. 4, bottom panel). It is worth noticing that the AT8 antibody did not stain for tau in axons in CA3 and DG.

**FIGURE 4.**
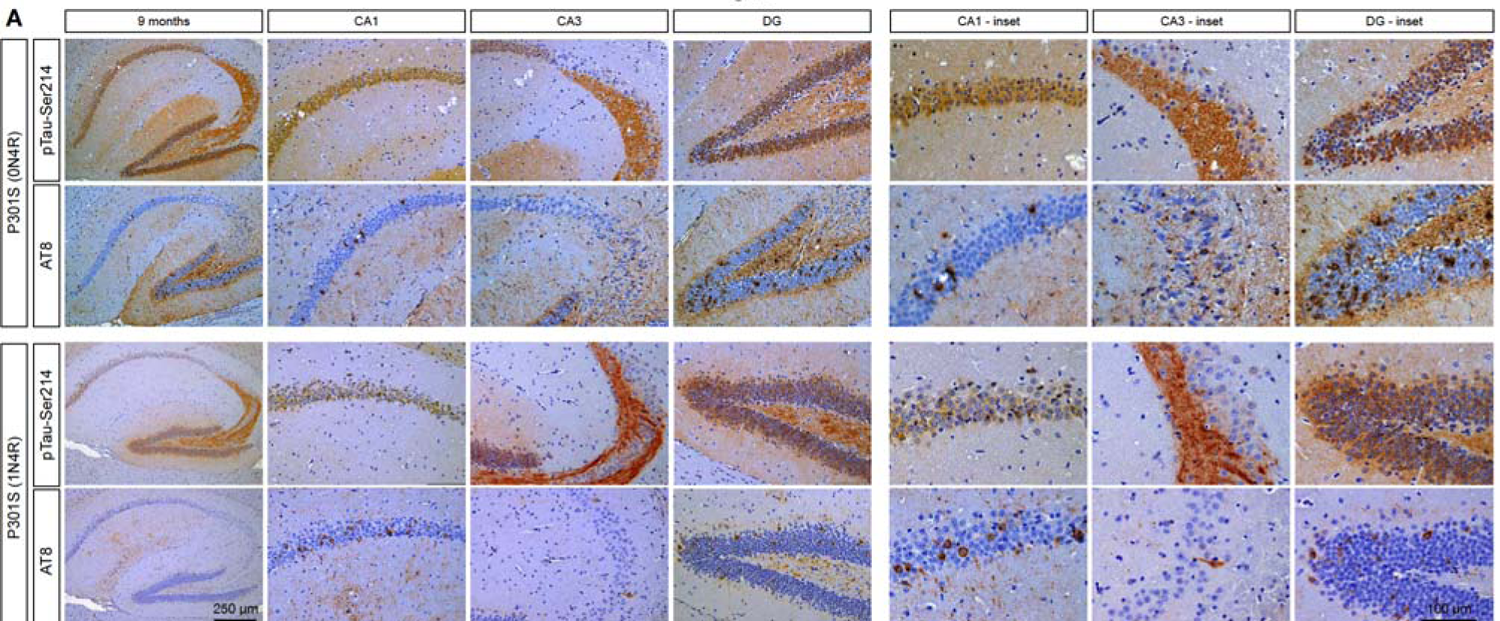
The staining pattern of tau phosphorylated at serine 214 differs from AT8-positive tau species in two different tauopathies mouse models. (**A**) Immunohistochemical staining of pTau-Ser214 and AT8 at 9 months old in both the P301S (0N4R, upper panel) and P301S (1N4R, lower panel) mouse models. Scale bar: 250 µm (low magnification hippocampus), and 100 µm (region-specific hippocampal insets); n=3.

To confirm this observation, we perform double immunofluorescences of pTau-Ser214 and AT8 in the hippocampal brain section of 9-month-old of both mouse models. Double immunostaining confirmed that the AT8 staining pattern differs from pTau-Ser214 in both models, where AT8 detected NFTs-like pathology and pTau-Ser214 resembled pre-tangles and tau in axons (Fig. 5A and 5B), suggesting the diversity of tau species within the brain.

**FIGURE 5.**
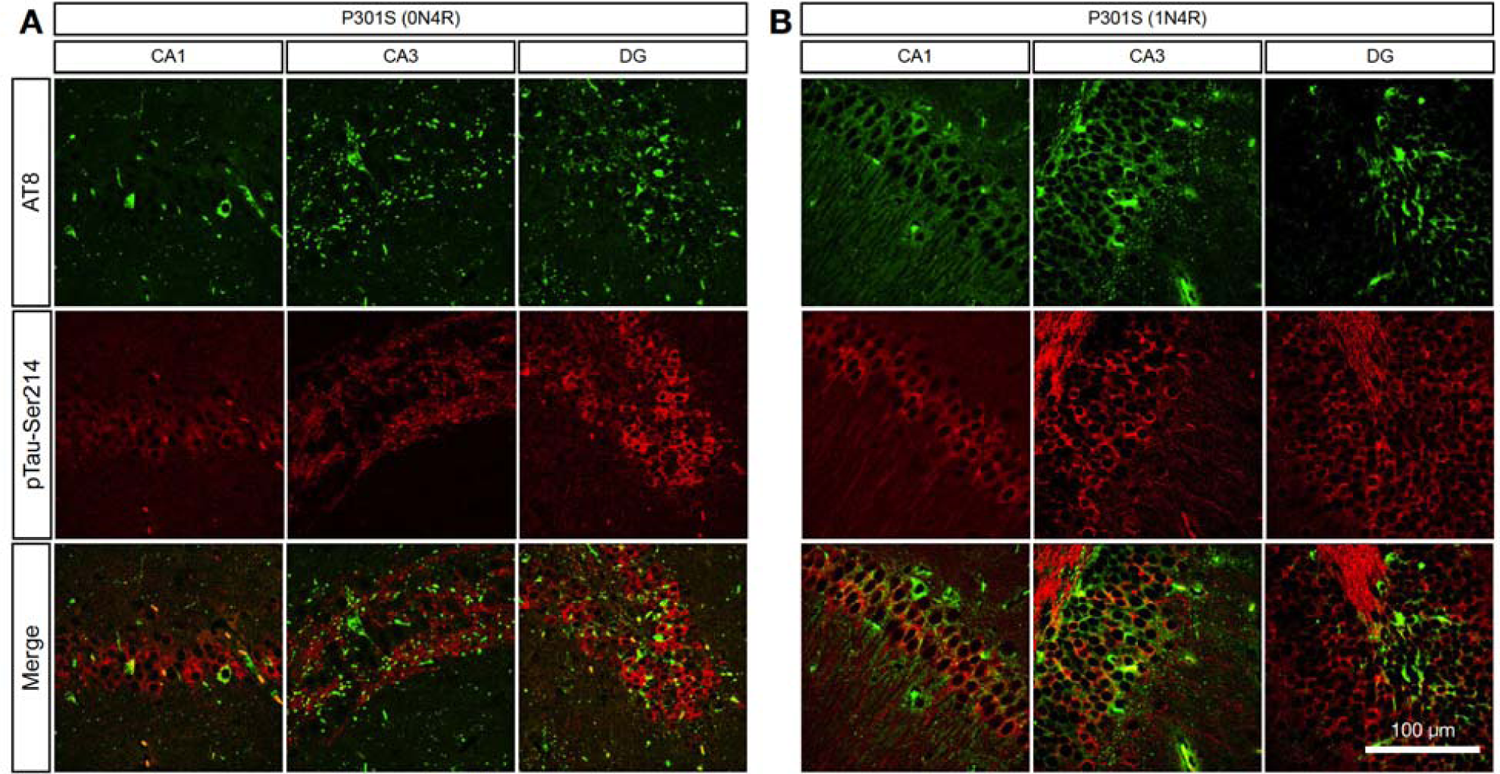
Tau phosphorylated at serine 214 is differentially distributed compared to AT8-positive staining. (**A, B**) Double immunofluorescence using specific antibodies against pTau-Ser214 (red) and AT8 (green) was performed in brain sections of 9-month-old brains in the (**A**) Tau P301S (0N4R) and (**B**) Tau P301S (1N4R) tauopathy models to analyze their spatial distribution in the hippocampus; n=3.

### Tau species phosphorylated at serine 214 but not at serine 202/ Threonine 205 (AT8) are deposited in the synaptic compartment

A recent study has demonstrated the presence of tau species with seeding activity in cytosolic and synaptic fractions from AD patients’ brain samples^17^. Based on this observation, we aimed to determine if there is a compartmentalized accumulation of pTau-Ser214 species with seeding activity present in the cytosol and/or synapses as observed in AD patients^17^. To do this, we isolated TBS-soluble synaptosomes and cytosolic fractions from the hippocampus of 9-month-old P301S (1N4R) animals (Fig. 6). Tau seeding activity in P301S (1N4R) mice was detected in both cytosolic and synaptic fractions (Fig. 6A). We then performed western blot analysis using anti pTau-Ser214, AT8, and HT7 tau antibodies; PSD95 was used to confirm synaptic compartment enrichment. The analysis revealed that pTau-Ser214 is accumulated in the cytosolic fraction and the synaptic compartment (Fig. 6B and 6E). Interestingly, AT8 is highly accumulated in the cytosolic fraction and was not detected in the synaptic fraction (Fig. 6C and 6E), with no changes in the distribution of total human tau quantified with HT7 antibody (Fig. 6D). Altogether, these results suggest differences in the accumulation of tau species between synaptic and cytosolic compartments, supporting the idea of the heterogeneity of pathological tau species with seeding activity that are not only within the same brain tissue but also within the same individual cell.

**FIGURE 6.**
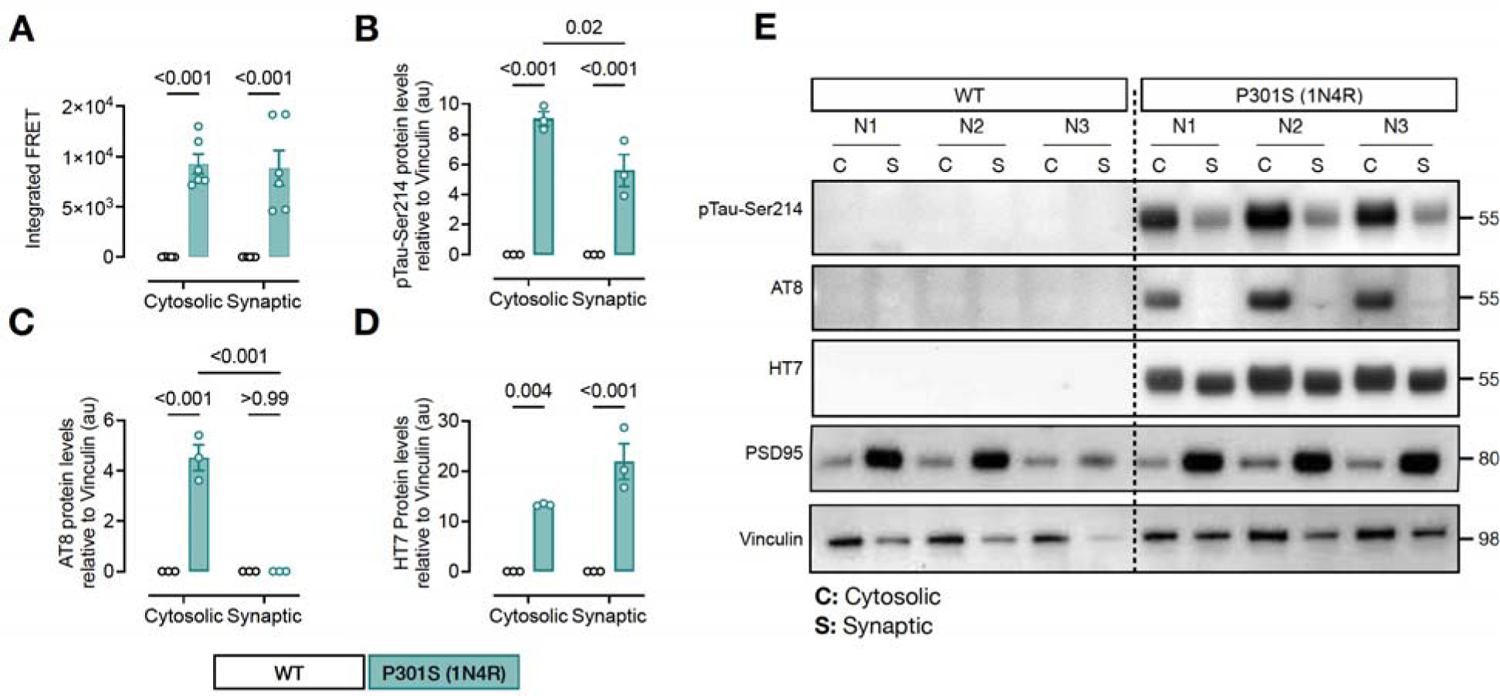
Tau phosphorylation at serine 214 is deposited in the synaptic compartment. **(A)** Tau seeding activity in cytosolic and synaptic TBS-soluble hippocampal fractions from 6 months old Tau P301S (1N4R) mouse model. Significance was determined by ordinary one-way ANOVA; n=6. (**B-E**) Quantifications of pTau-Ser214 **(B)**, AT8 **(C),** and HT7 **(D)** levels of a western blot (**E**) using cytosolic (**C**) and synaptic (**S**) fractions. PSD95 antibody was used to confirm synaptic compartment enrichment. Values are represented as the mean ± s.e.m, and significance was determined by two-way ANOVA; n=3.

### Phosphorylation of tau^P301L^ at serine 214 is involved in the transsynaptic propagation of pathological tau species

Considering previous studies suggesting that tau aggregates propagate transynaptically and our results showing that the levels of pTau-Ser214 correlate with tau seeding activity during disease progression and how this phospho-tau species accumulates at the synapse, we addressed the question of whether phosphorylation of tau at serine 214 has a pathological role in tau seeding activity as well as the transsynaptic propagation of tau. First, to evaluate the effect of pTau-Ser214 in seeding activity, we utilized a transgenic *Drosophila melanogaster* model developed in our lab, which consists of a fly overexpressing the human tau 2N4R carrying the P301L (tau^P301L^), or both P301L and S214A mutations (tau^P301L/S214A^) under the GMR-Gal4 driver (Fig. 7A). This *in vivo* model allows us to study the effect of introducing specific mutation to tau models in the context of the seeding activity and other pathological events as tau accumulation. We used head TBS-soluble lysate from 10-day-old flies to further quantify seeding activity in the HEK biosensor cell line. The overexpression of the tau^P301L^ mutant showed a robust seeding activity (Fig. 7B). Interestingly, when introducing the phospho-dead mutation S214A in tau^P301L^ flies, fly head lysates showed a slight but significant decrease in seeding activity (p=0.0223), suggesting that S214A may influence seeding activity but is not critical for tau-seed formation itself (Fig. 7B). Quantification of tau levels by western blot revealed that tau^P301L^, and tau^P301L/S214A^ flies had no significant differences on the accumulation of human tau (Fig. 7C).

**FIGURE 7.**
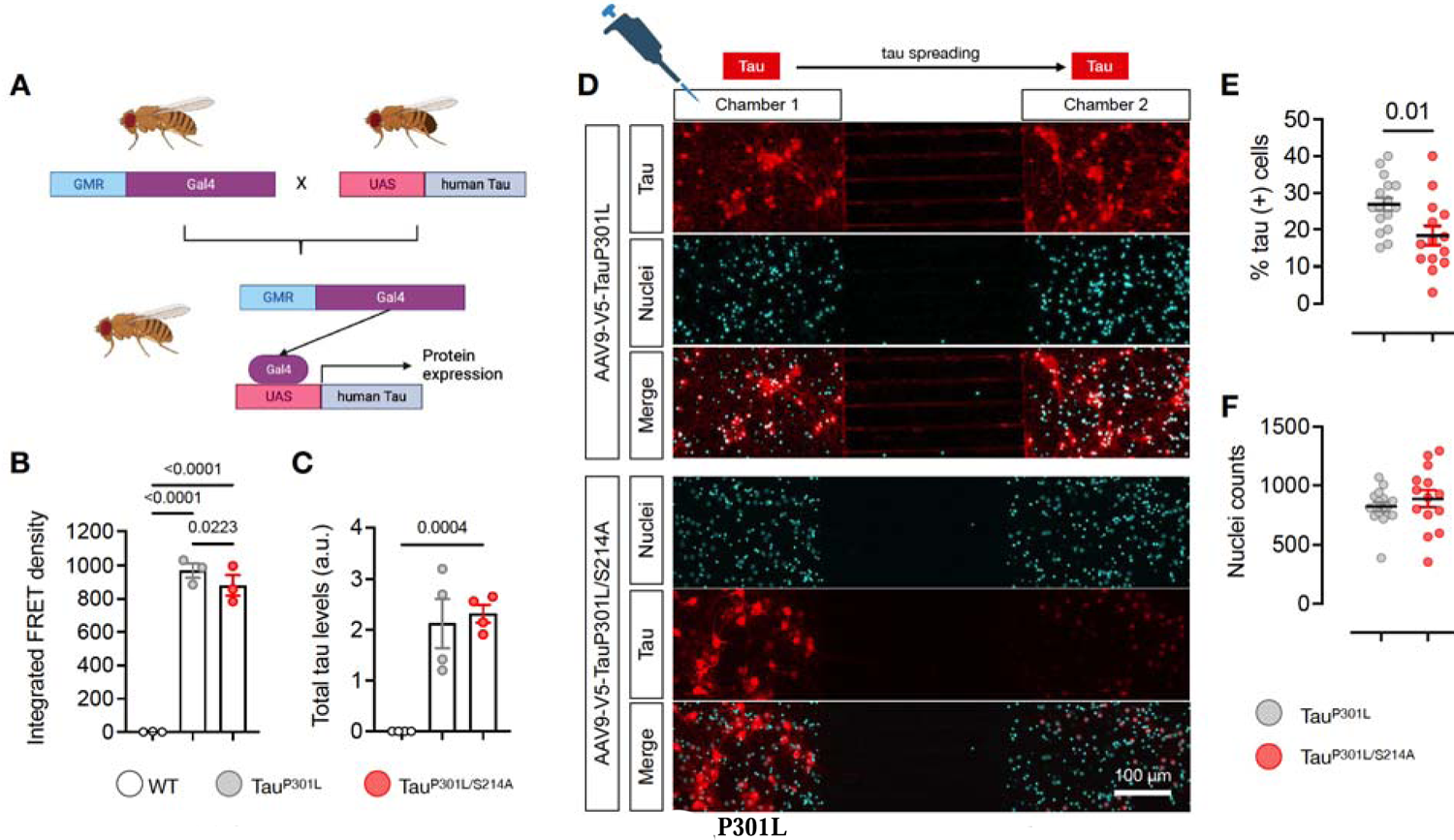
Phosphorylation of tau^P301L^ at serine 214 is involved in spreading pathological tau. **(A)** Schematic of the Gal4-UAS crosses utilized to overexpress human tau^P301^ or tau^P301L/S2^^14^ under the GMR driver (eye promoter). (**B**) Integrated FRET density (seeding activity) in fly head lysates from WT controls, and flies overexpressing human tau^P301L^ or tau^P301L/S214A^ mutations. Each dot represents a pool of 20 heads (n=20 with an equal number of males and females). (**C**) Total tau levels were quantified by western blot using the HT7 antibody to specifically detect human tau. Each dot represents a pool of 20 heads (n=20 an equal number of males and females). **(D)** Immunofluorescence against hTau (HT7, Red) in primary mouse neonatal cortical neurons plated in both microfluidic chambers. Neurons were transduced in the first chamber at 1 DIV with an AAV9 harboring plasmids coding for either tau^P301L^ or tau^P301L/S214A^ and tagged with V5. The second chamber shows the spreading of tau from the first chamber. Scale bar: 100 µm. **(E)** Quantification of Tau spreading in Chamber 2 and normalized by the number of nuclei in (**F**). Values are represented as the mean ± s.e.m from 3 independent experiments, and significance was determined by t-test.

In another set of experiments, to evaluate if phosphorylation of tau at serine 214 is relevant for transsynaptic propagation, we cultured mouse wild-type neonatal cortical neurons in microfluidic chambers and transduced them with two different AAV9, one coding for the human tau harboring the pathological mutation tau^P301L^, and another one harboring the tau^P301L/S214A^ double mutant. The microfluidic chamber facilitates the culture of neurons in two distinct chambers, which are separated by microchannels, allowing the visualization and quantification of the propagation of pathological tau from one chamber to the next one, as previously reported^40^. This design enables the extension of neuron axons through the microchannels, allowing them to establish synaptic connections with neurons located in the second chamber^41,42^. While one chamber was transduced with the tau encoding AAV9, the second chamber was not transduced, to allow tau to spread from the first chamber to the second one. Immunostainings against HT7 revealed that tau^P301L^ propagation was increased in the neurons in the second chamber (Fig. 7D-F), consistent with previous literature^40,43^. Interestingly, the replacement of the serine 214 by alanine prevented phosphorylation at this site and more importantly, significantly decreased the spreading of the pathological tau into the second chamber (p=0.01), supporting the notion of the role of pTau-Ser214 in tau transsynaptic spreading (Fig. 7D-F). Overall, these results suggest that the phosphorylation of tau at serine 214 is not an essential phosphorylation to promote the formation of the tau-seed but rather is important for tau to reach the synaptic terminal, facilitating the synaptic accumulation of tau which could be a key event for the transsynaptic propagation of the tau-seed.

## DISCUSSION

The accumulation and deposition of abnormal tau protein in the brain is a hallmark feature of tauopathies, a group of neurodegenerative disorders that includes Alzheimer’s disease. One important aspect of tau pathology is the spreading and seeding activity of these aberrant and toxic forms of tau, which refers to its ability to induce the aggregation of other tau molecules and promote the spread of tau pathology throughout the brain. Despite the significant progress in understanding tau pathology^17^, the precise cellular mechanism underlying tau propagation and the identity of the specific tau species involved in the seeding process remain unanswered. Our results support previous findings demonstrating tau seeding activity as early as 2 months in two mouse models of tauopathy, a significant period before the onset of synaptic impairments, glial activation, NFT pathology, and behavioral deficits^33,34^. Additionally, we found that the early seeding activity increases with age in these tau models. These observations suggest that tau seeding activity may represent an early and important step in the pathogenesis and temporal progression of tauopathy and that targeting this process may represent a promising therapeutic strategy.

Toxic aberrant tau species can be found in a hyperphosphorylated and aggregated state leading to the formation of NFTs, which form insoluble fibrils that accumulate within the neurons, leading to dysfunction and eventual death^44,45^. Nevertheless, a growing literature on tauopathies suggests that NFTs are not the primary toxic entities in these disorders^33,46–48^. Instead, accumulating evidence suggests that soluble hyperphosphorylated tau species are the primary mediators of seeding activity and neurotoxicity^33,49–53^. In addition, it has been observed that different tauopathies exhibit a great diversity of hyperphosphorylated tau aggregates, which correspond to distinct pathological tau “strains” that are associated with different clinical manifestations^24,27^. The identification of these pathological tau strains adds a layer of complexity when attempting to block the spread of tau seed, which has proven difficult due to the unknown structure of the tau seed and the recent studies confirming the molecular diversity of seeding-competent tau in both in vitro and in vivo models^23–27^. This complexity makes it exceedingly challenging to determine the exact epitope or post-translational modification to target tau. Here, we performed a detailed correlation analysis between a diverse set of phospho-tau and conformational tau species with seeding activity and demonstrated that the tau phosphorylation at serine 214 (pTau-Ser214) in the only one that positively correlated with seeding activity in two different mouse models of tauopathy during disease progression. This suggests that the pTau-Ser214 site represents a possible candidate as a reliable biomarker to track and correlate the evolution and progression of tau species with seeding capability, within individuals and animal models of tauopathy.

Interestingly, at an early stage of the disease, we observed that pTau-Ser214 has a distinct histopathological distribution from other tau species that are positive for the AT8 antibody, a marker for late-stage tau pathology. AT8 has been widely utilized for tau pathology detection, particularly in the form of NFTs by IHC in AD patients^12,42^. Its relevance lies in its ability to identify and quantify the accumulation of NFT in brain tissue samples, providing a tool to determine the AD stage. Our results demonstrate that AT8-positive tau species are absent in synaptic compartments. On the other hand, we found that pTau-Ser214 is deposited in the synaptic compartment. This is an important observation, not only because the synaptic compartment is a critical site for neuronal communication and function, but also because the deposition of hyperphosphorylated tau species at this site may contribute to the cognitive and behavioral deficits observed in tauopathies and could promote the transsynaptic propagation of tau^17^. Synaptic accumulation of pTau-Ser214 with seeding activity and the distinct pattern to AT8 species not only reveals the diversity in aberrant tau conformations and phosphorylation states within the brain during disease progression but also supports the idea that phosphorylation at serine 214 could be key for toxic tau species to reach and accumulate at the synapse and subsequently trigger the transsynaptic propagation. Recent work from our lab established a correlation between behavioral deficits and different pathological tau species in a mouse model for tauopathy, suggesting that each tau species analyzed (including soluble pTau-Ser214) could be responsible for a variety of behavioral deficits in PS19 (1N4R tau isoform) mouse model^35^. Other studies suggest that the structural diversity of tau aggregates may contribute to the differences in symptoms and pathologies observed in neurodegenerative tauopathies^54^. However, conclusive mechanistic investigations of tau proteins face significant barriers due to the inconsistent tau species being studied, which is crucial since the aggregated state of tau plays a critical role in its pathological evolution^55^. Moreover, tau can exhibit conformational differences, leading to various downstream effects^27,56^, based on a novel study showing that different tau conformations resulting from the Tau^P301L^ mutation can drive distinct phenotypes in FTLD patients^57^.

The phosphorylation of tau at specific sites can influence other toxic characteristics. For instance, phosphorylation of tau at specific sites has been shown to influence tau aggregation and tau seeding activity. Similarly, phosphorylation at specific sites may contribute to a greater propensity of tau to spread/propagate and the pattern in which propagates^58,59^. Our result suggested that pTau-Ser214 is involved in the transsynaptic propagation of pathological tau in vitro and contributes little to in vivo seeding activity. Taking into consideration that pTau-Ser214 is also enriched in the synaptic terminal, altogether these results suggest that tau phosphorylation at serine 214 is not required for the formation of the tau-seed but rather is essential to reach the synaptic terminal, therefore facilitating tau transsynaptic transmission.

In conclusion, our study establishes the heterogeneity of tau species with seeding activity not only in the same brain region but at the cellular level as well. Additionally, our findings reveal that tau phosphorylation at serine 214 is an early event in tau pathology, and its levels strongly correlate with seeding activity in an age-dependent manner in two mouse models for tauopathies. Even more, our results suggest that phosphorylation of tau at serine 214 does not significantly affect the formation of the tau-seed but is essential for the accumulation of tau at the synapses which is a key event for the transsynaptic propagation of tau aggregates. Altogether, this study supports the importance of precision strategies to develop tau biomarkers to measure disease progression during aging tailored to the individual patient.

## DATA AVAILABILITY

Source data are provided in this paper. All other numerical data are available from the corresponding author upon request.

## FUNDING

This work was supported by NIH/NINDS 1R01NS119280, NIH/NIA 1R01AG059639, NIH/NIA R21AG075541grants and ALZDISCOVERY-1049108 from the Alzheimer’s Association to C.A.L.-R., AARFD-21-847663 to P.M. This publication was also supported by the Sara Roush Memorial Fellowship in Alzheimer’s Disease, the Indiana Alzheimer’s Disease Research Center, and the Stark Neurosciences Research Institute, and made possible by the Indiana Clinical and Translational Sciences Institute, funded in part by grant # UL1TR002529 from the National Institutes of Health, National Center for Advancing Translational Sciences. The funders had no role in the study design, data collection and analysis, decision to publish, or manuscript preparation.

## AUTHORS CONTRIBUTIONS

P.M. and C.A.L.-R. conceived and designed the project with contributions from A.P., N.J-G., H.P., Y.Y., YJ. Y., and A.L.-G., performing experiments. A.P., Y.Y., and YJ. Y performed the correlation study using western blots for all tau antibodies and FRET assays. A.L.-G. helped with AT8 and S214 IHQ. P.M., and H.P. performed IF for AT8 and S214 characterization. N.J-G. performed Synaptic and Cytosolic isolation and characterization. P.M. designed and performed microfluidic experiments. R.V. provided mice and helped with proofreading. Funding acquisition from: C.A.L.-R and P.M. P.M. and C.A.L.-R wrote the manuscript with contributions from all the authors.

## COMPETING INTEREST

The authors declare no competing interest.

## Notes

### Competing Interest Statement

The authors have declared no competing interest.

## REFERENCES

1. Johnson GVW, Stoothoff WH. Tau phosphorylation in neuronal cell function and dysfunction. J Cell Sci. 2004;117(24):5721–5729. doi:10.1242/jcs.01558

2. Liu F, Gong CX. Tau exon 10 alternative splicing and tauopathies. Mol Neurodegener. 2008;3(1):8. doi:10.1186/1750-1326-3-8

3. Goedert M, Spillantini MG, Jakes R, Rutherford D, Crowther RA. Multiple isoforms of human microtubule-associated protein tau: sequences and localization in neurofibrillary tangles of Alzheimer’s disease. Neuron. 1989;3(4):519–526. doi:10.1016/0896-6273(89)90210-9

4. Gendron TF, Petrucelli L. The role of tau in neurodegeneration. Mol Neurodegener. 2009;4(1):13. doi:10.1186/1750-1326-4-13

5. Arriagada PV, Growdon JH, Hedley-Whyte ET, Hyman BT. Neurofibrillary tangles but not senile plaques parallel duration and severity of Alzheimer’s disease. Neurology. 1992;42(3 Pt 1):631–639. doi:10.1212/wnl.42.3.631

6. Gómez Isla T, Hollister R, West H, et al. Neuronal loss correlates with but exceeds neurofibrillary tangles in Alzheimer’s disease. Ann Neurol. 1997;41(1):17–24. doi:10.1002/ana.410410106

7. Augustinack JC, Schneider A, Mandelkow EM, Hyman BT. Specific tau phosphorylation sites correlate with severity of neuronal cytopathology in Alzheimer’s disease. Acta Neuropathol. 2002;103(1):26–35. doi:10.1007/s004010100423

8. Steinhilb ML, Dias-Santagata D, Fulga TA, Felch DL, Feany MB. Tau Phosphorylation Sites Work in Concert to Promote Neurotoxicity In Vivo. Mol Biol Cell. 2007;18(12):5060–5068. doi:10.1091/mbc.e07-04-0327

9. Hong M, Lee VMY. Insulin and Insulin-like Growth Factor-1 Regulate Tau Phosphorylation in Cultured Human Neurons*. J Biol Chem. 1997;272(31):19547–19553. doi:10.1074/jbc.272.31.19547

10. Drechsel DN, Hyman AA, Cobb MH, Kirschner MW. Modulation of the dynamic instability of tubulin assembly by the microtubule-associated protein tau. Mol Biol Cell. 1992;3(10):1141–1154. doi:10.1091/mbc.3.10.1141

11. Xia Y, Prokop S, Giasson BI. “Don’t Phos Over Tau”: recent developments in clinical biomarkers and therapies targeting tau phosphorylation in Alzheimer’s disease and other tauopathies. Mol Neurodegener. 2021;16(1):37. doi:10.1186/s13024-021-00460-5

12. Braak H, Braak E. Frequency of Stages of Alzheimer-Related Lesions in Different Age Categories. Neurobiol Aging. 1997;18(4):351–357. doi:10.1016/s0197-4580(97)00056-0

13. Rodriguez RD, Grinberg LT. Argyrophilic grain disease: An underestimated tauopathy. Dementia Neuropsychologia. 2014;9(1):2–8. doi:10.1590/s1980-57642015dn91000002

14. Murray ME, DeTure M. APA handbook of dementia. Published online 2018:41–66. doi:10.1037/0000076-003

15. Chung Deun C, Roemer S, Petrucelli L, Dickson DW. Cellular and pathological heterogeneity of primary tauopathies. Mol Neurodegener. 2021;16(1):57. doi:10.1186/s13024-021-00476-x

16. Fuster-Matanzo A, Hernández F, Ávila J. Tau Spreading Mechanisms; Implications for Dysfunctional Tauopathies. Int J Mol Sci. 2018;19(3):645. doi:10.3390/ijms19030645

17. DeVos SL, Corjuc BT, Oakley DH, et al. Synaptic Tau Seeding Precedes Tau Pathology in Human Alzheimer’s Disease Brain. Front Neurosci. 2018;12:267. doi:10.3389/fnins.2018.00267

18. Wu JW, Hussaini SA, Bastille IM, et al. Neuronal activity enhances tau propagation and tau pathology in vivo. Nat Neurosci. 2016;19(8):1085–1092. doi:10.1038/nn.4328

19. de Calignon A, Polydoro M, Suárez-Calvet M, et al. Propagation of Tau Pathology in a Model of Early Alzheimer’s Disease. Neuron. 2012;73(4):685–697. doi:10.1016/j.neuron.2011.11.033

20. Lasagna-Reeves CA, Castillo-Carranza DL, Sengupta U, et al. Alzheimer brain-derived tau oligomers propagate pathology from endogenous tau. Sci Rep-uk. 2012;2(1):700. doi:10.1038/srep00700

21. Tanaka Y, Yamada K, Satake K, et al. Seeding Activity-Based Detection Uncovers the Different Release Mechanisms of Seed-Competent Tau Versus Inert Tau via Lysosomal Exocytosis. Front Neurosci. 2019;13:1258. doi:10.3389/fnins.2019.01258

22. Guo JL, Narasimhan S, Changolkar L, et al. Unique pathological tau conformers from Alzheimer’s brains transmit tau pathology in nontransgenic mice. J Exp Medicine. 2016;213(12):2635–2654. doi:10.1084/jem.20160833

23. Narasimhan S, Guo JL, Changolkar L, et al. Pathological Tau Strains from Human Brains Recapitulate the Diversity of Tauopathies in Nontransgenic Mouse Brain. J Neurosci. 2017;37(47):11406–11423. doi:10.1523/jneurosci.1230-17.2017

24. Clavaguera F, Akatsu H, Fraser G, et al. Brain homogenates from human tauopathies induce tau inclusions in mouse brain. Proc National Acad Sci. 2013;110(23):9535–9540. doi:10.1073/pnas.1301175110

25. Kaufman SK, Diamond MI. Prion-Like Propagation of Protein Aggregation and Related Therapeutic Strategies. Neurotherapeutics. 2013;10(3):371–382. doi:10.1007/s13311-013-0196-3

26. Kaufman SK, Sanders DW, Thomas TL, et al. Tau Prion Strains Dictate Patterns of Cell Pathology, Progression Rate, and Regional Vulnerability In Vivo. Neuron. 2016;92(4):796–812. doi:10.1016/j.neuron.2016.09.055

27. Sanders DW, Kaufman SK, DeVos SL, et al. Distinct Tau Prion Strains Propagate in Cells and Mice and Define Different Tauopathies. Neuron. 2014;82(6):1271–1288. doi:10.1016/j.neuron.2014.04.047

28. Albert M, Mairet-Coello G, Danis C, et al. Prevention of tau seeding and propagation by immunotherapy with a central tau epitope antibody. Brain. 2019;142(6):1736–1750. doi:10.1093/brain/awz100

29. Yanamandra K, Jiang H, Mahan TE, et al. Anti-tau antibody reduces insoluble tau and decreases brain atrophy. Ann Clin Transl Neur. 2015;2(3):278–288. doi:10.1002/acn3.176

30. Courade JP, Angers R, Mairet-Coello G, et al. Epitope determines efficacy of therapeutic anti-Tau antibodies in a functional assay with human Alzheimer Tau. Acta Neuropathol. 2018;136(5):729–745. doi:10.1007/s00401-018-1911-2

31. Dujardin S, Commins C, Lathuiliere A, et al. Tau molecular diversity contributes to clinical heterogeneity in Alzheimer’s disease. Nat Med. 2020;26(8):1256–1263. doi:10.1038/s41591-020-0938-9

32. Holmes BB, Furman JL, Mahan TE, et al. Proteopathic tau seeding predicts tauopathy in vivo. Proc National Acad Sci. 2014;111(41):E4376–E4385. doi:10.1073/pnas.1411649111

33. Yoshiyama Y, Higuchi M, Zhang B, et al. Synapse Loss and Microglial Activation Precede Tangles in a P301S Tauopathy Mouse Model. Neuron. 2007;53(3):337–351. doi:10.1016/j.neuron.2007.01.010

34. Garringer HJ, Murrell J, Sammeta N, Gnezda A, Ghetti B, Vidal R. Increased Tau Phosphorylation and Tau Truncation, and Decreased Synaptophysin Levels in Mutant BRI2/Tau Transgenic Mice. Plos One. 2013;8(2):e56426. doi:10.1371/journal.pone.0056426

35. Patel H, Martinez P, Perkins A, et al. Pathological tau and reactive astrogliosis are associated with distinct functional deficits in a mouse model of tauopathy. Neurobiol Aging. 2022;109:52–63. doi:10.1016/j.neurobiolaging.2021.09.006

36. Martinez P, Patel H, You Y, et al. Bassoon contributes to tau-seed propagation and neurotoxicity. Nat Neurosci. 2022;25(12):1597–1607. doi:10.1038/s41593-022-01191-6

37. Southam KA, King AE, Blizzard CA, McCormack GH, Dickson TC. Microfluidic primary culture model of the lower motor neuron–neuromuscular junction circuit. J Neurosci Meth. 2013;218(2):164–169. doi:10.1016/j.jneumeth.2013.06.002

38. Cook C, Kang SS, Carlomagno Y, et al. Tau deposition drives neuropathological, inflammatory and behavioral abnormalities independently of neuronal loss in a novel mouse model. Hum Mol Genet. 2015;24(21):6198–6212. doi:10.1093/hmg/ddv336

39. Mercken M, Vandermeeren M, Lübke U, et al. Monoclonal antibodies with selective specificity for Alzheimer Tau are directed against phosphatase-sensitive epitopes. Acta Neuropathol. 1992;84(3):265–272. doi:10.1007/bf00227819

40. Katsikoudi A, Ficulle E, Cavallini A, et al. Quantitative propagation of assembled human Tau from Alzheimer’s disease brain in microfluidic neuronal cultures. J Biol Chem. 2020;295(37):13079–13093. doi:10.1074/jbc.ra120.013325

41. Bomba-Warczak E, Vevea JD, Brittain JM, et al. Interneuronal Transfer and Distal Action of Tetanus Toxin and Botulinum Neurotoxins A and D in Central Neurons. Cell Reports. 2016;16(7):1974–1987. doi:10.1016/j.celrep.2016.06.104

42. Taylor AM, Blurton-Jones M, Rhee SW, Cribbs DH, Cotman CW, Jeon NL. A microfluidic culture platform for CNS axonal injury, regeneration and transport. Nat Methods. 2005;2(8):599–605. doi:10.1038/nmeth777

43. Dujardin S, Lécolle K, Caillierez R, et al. Neuron-to-neuron wild-type Tau protein transfer through a trans-synaptic mechanism: relevance to sporadic tauopathies. Acta Neuropathologica Commun. 2014;2(1):14. doi:10.1186/2051-5960-2-14

44. Buée L, Troquier L, Burnouf S, et al. From tau phosphorylation to tau aggregation: what about neuronal death? Biochem Soc T. 2010;38(4):967–972. doi:10.1042/bst0380967

45. Chun W, Johnson GVW. The role of tau phosphorylation and cleavage in neuronal cell death. Frontiers Biosci J Virtual Libr. 2006;12(1):733–756. doi:10.2741/2097

46. Berger Z, Roder H, Hanna A, et al. Accumulation of Pathological Tau Species and Memory Loss in a Conditional Model of Tauopathy. J Neurosci. 2007;27(14):3650–3662. doi:10.1523/jneurosci.0587-07.2007

47. Castillo-Carranza DL, Sengupta U, Guerrero-Muñoz MJ, et al. Passive Immunization with Tau Oligomer Monoclonal Antibody Reverses Tauopathy Phenotypes without Affecting Hyperphosphorylated Neurofibrillary Tangles. J Neurosci. 2014;34(12):4260–4272. doi:10.1523/jneurosci.3192-13.2014

48. SantaCruz K, Lewis J, Spires T, et al. Tau Suppression in a Neurodegenerative Mouse Model Improves Memory Function. Science. 2005;309(5733):476–481. doi:10.1126/science.1113694

49. Haroutunian V, Davies P, Vianna C, Buxbaum JD, Purohit DP. Tau protein abnormalities associated with the progression of alzheimer disease type dementia. Neurobiol Aging. 2007;28(1):1–7. doi:10.1016/j.neurobiolaging.2005.11.001

50. Lasagna-Reeves CA, Castillo-Carranza DL, Sengupta U, Clos AL, Jackson GR, Kayed R. Tau oligomers impair memory and induce synaptic and mitochondrial dysfunction in wild-type mice. Mol Neurodegener. 2011;6(1):39. doi:10.1186/1750-1326-6-39

51. Lasagna Reeves CA, Castillo Carranza DL, Sengupta U, et al. Identification of oligomers at early stages of tau aggregation in Alzheimer’s disease. Faseb J. 2012;26(5):1946–1959. doi:10.1096/fj.11-199851

52. Maeda S, Takashima A. Tau Biology. Adv Exp Med Biol. 2020;1184:373–380. doi:10.1007/978-981-32-9358-8_27

53. Takashima A. Hyperphosphorylated Tau is a Cause of Neuronal Dysfunction in Tauopathy. J Alzheimer’s Dis. 2008;14(4):371–375. doi:10.3233/jad-2008-14403

54. Gerson JE, Mudher A, Kayed R. Potential mechanisms and implications for the formation of tau oligomeric strains. Crit Rev Biochem Mol. 2016;51(6):482–496. doi:10.1080/10409238.2016.1226251

55. Cowan CM, Mudher A. Are Tau Aggregates Toxic or Protective in Tauopathies? Front Neurol. 2013;4:114. doi:10.3389/fneur.2013.00114

56. Hyman BT. Tau Propagation, Different Tau Phenotypes, and Prion-like Properties of Tau. Neuron. 2014;82(6):1189–1190. doi:10.1016/j.neuron.2014.06.004

57. Daude N, Kim C, Kang SG, et al. Diverse, evolving conformer populations drive distinct phenotypes in frontotemporal lobar degeneration caused by the same MAPT-P301L mutation. Acta Neuropathol. 2020;139(6):1045–1070. doi:10.1007/s00401-020-02148-4

58. Brunello CA, Merezhko M, Uronen RL, Huttunen HJ. Mechanisms of secretion and spreading of pathological tau protein. Cell Mol Life Sci. 2020;77(9):1721–1744. doi:10.1007/s00018-019-03349-1

59. Dujardin S, Bégard S, Caillierez R, et al. Different tau species lead to heterogeneous tau pathology propagation and misfolding. Acta Neuropathologica Commun. 2018;6(1):132. doi:10.1186/s40478-018-0637-7

